## Supplementary material for "Maternal behavior influences vocal practice and learning processes in the greater sac-winged bat"

### Supplemental Information

**Table S1. Ethogram of behaviors during babbling bouts.**

| Behavior | Description | Observed in: |
|---|---|---|
| crawl away* | One individual crawls a certain distance (> 2 cm) away from the other. | P / M |
| crawl towards* | One individual crawls towards another individual (approaches up to 2 cm). | P / M |
| hover* | Hover flight in front of an individual (for a few seconds). | P / M |
| wing-poke* | Poking a conspecific with wrist. Pups normally repeatedly poke their mothers. Mothers usually poke the pup only once to terminate the babbling bout. Includes wing pokes with and without physical contact. | P / M |
| short flight* | Individual flies to another spot in the day roost or individual flies towards interaction partner. | P / M |
| rock | Rocking with entire body from side to side; also used when mother wants pup to detach from teat. | P / M |
| wing stretch | Wing is stretched out completely. | P / M |
| head stretch | Head and neck are conspicuously bent and stretched to either left or right side. | P |
| push up | Push wrists against wall and push body up. | P / M |
| dribble | Pound with wrists against wall, very fast and repetitive. | P |
| wrist lift | Pound with wrists against wall once, not hitting other individual. | P |
| babble | Vocal practice. | P |

The behaviors labelled with a * either occurred as solitary behavior or as behavioral sequence (i.e. interaction). These interactive behaviors were used to assess the maternal behavioral influence on pup babbling. The behaviors without a * (with the exception of babble) were defined as comfort behaviors and not included as behavioral feedback. The third column depicts if the behavior was observed in the pup (P) or both pup and mother (P/M).

**Table S2. Maternal display rates: duration of babbling is not a confounding factor**

| Response variable | Parameter | Estimate (SE) | Significance |
|---|---|---|---|
| **A** *Babbling bout duration [s]* | *Fixed effects* | | |
| | Intercept | 6.132 (0.091) | p < 0.001 |
| | Mat. behav. rates during babbling (standardized) | -0.146 (0.053) | **p = 0.006** |
| | Pup age (standardized) | 0.418 (0.050) | **p < 0.001** |
| | No. of tutors (standardized) | -0.063 (0.095) | p = 0.51 |
| | *Random effect (standard deviation)* | | |
| | Pup ID | 0.24 | |

The number of displays in longer bouts could just reflect that more displays are possible in a longer period. To assess this potential confounding factor, we investigated the effect of display rates on vocal bout duration (N=19 pups, N=186 babbling bouts, GLMM family Gamma with log link).

### Maternal influence on the amount of vocal practice.

#### *Maternal behaviors*

The number of analyzed bouts did not bias the analyzed maternal behavioral response (Spearman-rank correlation test: p = 0.8, rho = -0.07, N = 19 pups). The number of maternal behaviors did not differ between pup sex (Mann-Whitney-U-Test: z = -0.063888, p = 1, N = 14 pups).

#### *Bout duration*

Bout duration did not differ between pup sex (Mann-Whitney-U-Test: z = -0.44721, p = 0.7, N=14 pups) or between colonies (Kruskal-Wallis Test: chi-squared = 7.6853, p = 0.2, N = 19 pups). However, bout duration was different between the two study populations, Costa Rica and Panama, with longer mean bout durations found in Panama (Mann-Whitney-U-Test: z = -2.0412, p = 0.04, N = 19 pups, average bout durations: Costa Rica: ~ 7 min, Panama: ~ 10 min).

#### *Babbling phase duration*

Babbling phase duration did not differ between pup sex (Mann-Whitney-U-Test: z = -0.51279, p = 0.6, N = 14 pups) or between colonies (Kruskal-Wallis Test: chi-squared, 11,224, p = 0.047, N = 14 pups, followed by pairwise Wilcox. test with Bonferroni correction: ns). However, the babbling phase duration was different between the two populations, with longer babbling phases present in pups from Costa Rica (Mann-Whitney-U-Test: z = 2.5754, p = 0.009, N = 19 pups, average babbling phase duration: Costa Rica: 52.7 days, Panama: 36.3 days).

### The influence of the social environment on learned syllables.

#### *Total number of syllables within babbling bouts*

Female and male pups did not differ in the number of buzz syllables they produced during babbling (Mann-Whitney-U-Test: z = 0.35355, p = 0.86, N = 7 pups). Likewise, there was no significant difference in the amount of buzz syllables produced between the different colonies (Kruskal-Wallis-Test: chi-squared = 2.0218, p = 0.37, N = 10 pups).

#### *Syllable versatility*

There was no difference between the syllabic diversity between male and female pups (Mann-Whitney-U-Test: z = -1.4142, p = 0.23, N = 7 pups) or between colonies (Kruskal-Wallis-Test: chi-squared = 0.54909, p = 0.76, N = 10 pups).

**Table S3. The average proportion of syllable type production.**

| ID | B1 | B2 | B2-trill | B3 | B4 |
|---|---|---|---|---|---|
| 11 | 0 | 47.8 | 6.6 | 24.7 | 12.6 |
| 12 | 25.8 | 56.1 | 6.6 | 9.7 | 1.8 |
| 13 | 0.13 | 50.2 | 12.7 | 16.1 | 6.7 |
| 14 | 0 | 54.9 | 25.8 | 27.8 | 19.2 |
| 15 | 0.1 | 54.9 | 16.2 | 9.0 | 19.7 |
| 16 | 4.9 | 67.2 | 1.6 | 20.6 | 5.7 |
| 17 | 2.3 | 73.5 | 3.4 | 20.4 | 0.4 |
| 18 | 0 | 54.2 | 14.5 | 24.1 | 7.2 |
| 19 | 0 | 53.7 | 7.8 | 24.8 | 13.8 |
| 20 | 0 | 50.9 | 24.5 | 11.6 | 5.3 |

For each pup, the average proportion of each learned syllable type produced during all analysed babbling bouts is displayed. Averaged over all pups B2 is the most commonly produced syllable type within babbling bouts (53.8%), followed by B3 (18.6%), B2 trill (11.4%), B4 (8.9%), and B1 (2.8%).

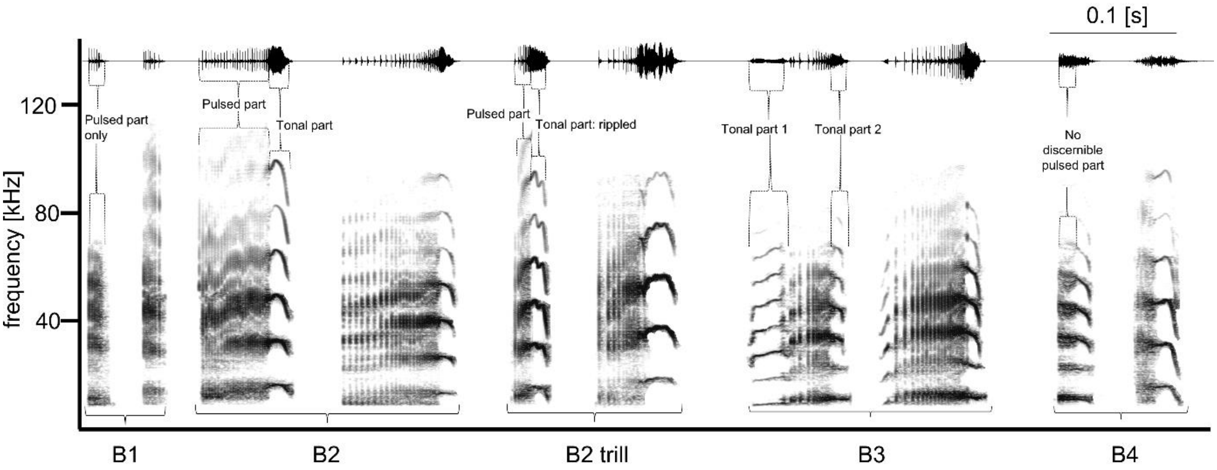


**Fig. S1. Five different song syllable types in babbling bouts.** We defined five different syllable types based on spectro-temporal characteristics (i.e. best visualized by the combination of the spectrogram and the oscillogram). B1 has no tonal part. B2 has a clear pulsed part connected to one tonal part. B2 trill has a pulsed part and a tonal part with rippled frequency modulations. B3 has at least two tonal parts connected to a pulsed part. B4 has no discernible pulsed part but only a smeared noisy part. These syllables can be flexibly used for the composition of territorial songs; their composition is usually an indication of the current aggression level of the singer.  The spectrogram was created in Avisoft SasLab Pro: Hamming window with 1024-point Fourier transformation and 87.5% overlap (frequency resolution: 244 Hz, time resolution: 0.512 ms).

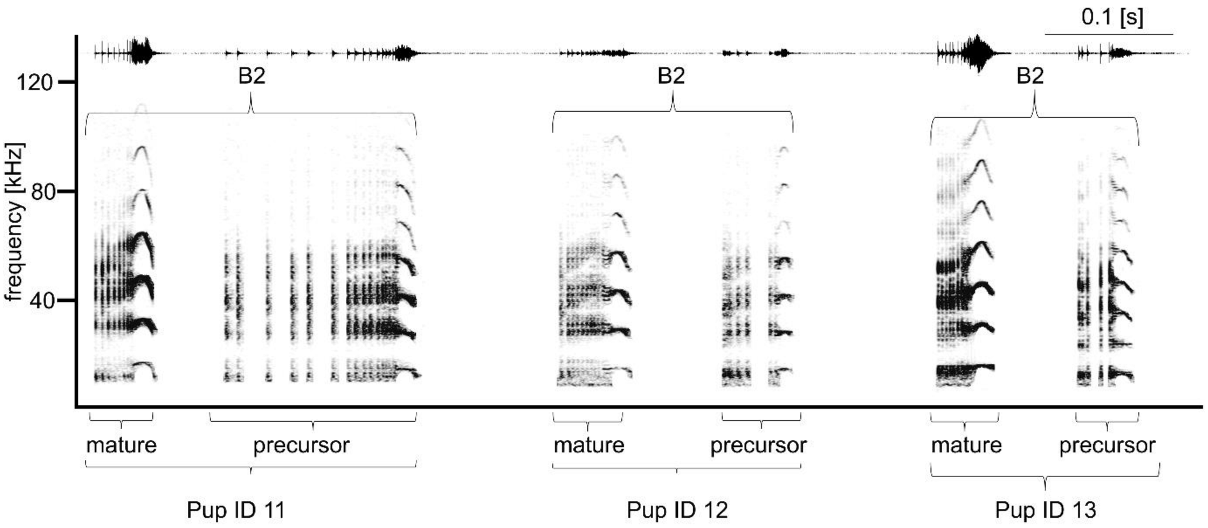


**Fig. S2. Precursor versus mature syllables.** This spectrogram illustrates the difference between precursor versus mature syllables (for the most common type B2 which was used in this analysis) from three different pups. Precursor syllables have distinct gaps, usually in the pulsed part, between the pulsed and the tonal part or even both. The spectrogram was created in Avisoft SASLab Pro: Hamming window with 1024-point Fourier transformation and 87.5% overlap (frequency resolution: 244 Hz, time resolution: 0.512 ms).

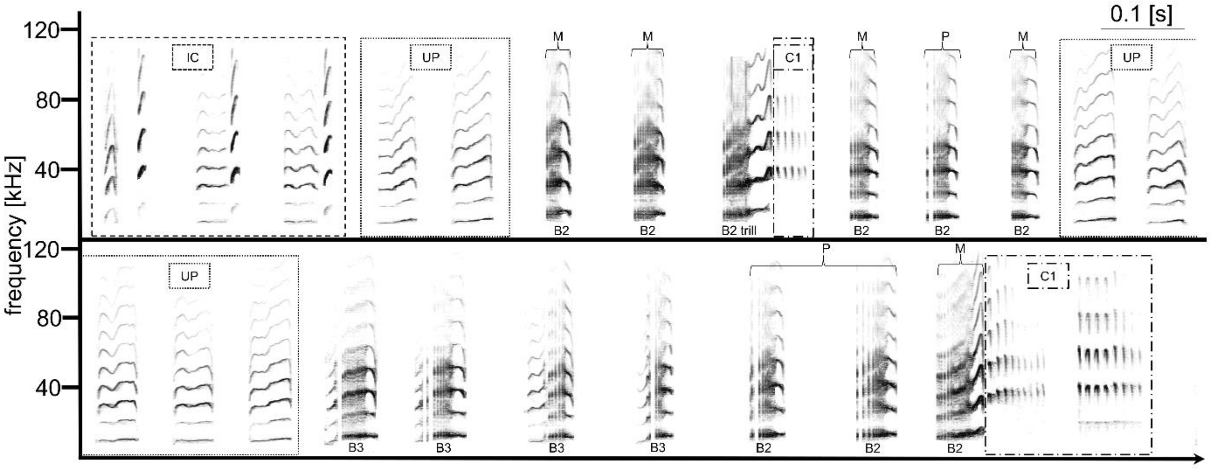


**Fig. S3. Babbling excerpt with syllable labeling.** This excerpt of a babbling bout (pup ID 13) illustrates the acoustic analysis which was performed in order to analyse a) the total number of song syllables (=amount of all song syllables; in this example the number of B2, B2 trill and B3), b) the versatility per bout (=amount of all different types per bout, subsequent calculation of the diversity index using the R package vegan), and, c) the number of precursor and mature B2 syllables within a babbling bout (= amount of mature (M) and precursor (P) B2 syllables). B2, B2 trill, and B3 belong to the adult-like syllables, being part of the later adult vocal repertoire. Besides the learned song syllables, babbling includes other adult-like syllable types (IC: Isolation call syllables, C1: chatter syllables) and syllables exclusively produced during babbling by pups (UP: undifferentiated proto-syllables). The spectrogram was created in Avisoft SASLab Pro: Hamming window with 1024-point Fourier transformation and 87.5% overlap (frequency resolution: 244 Hz, time resolution: 0.512 ms).

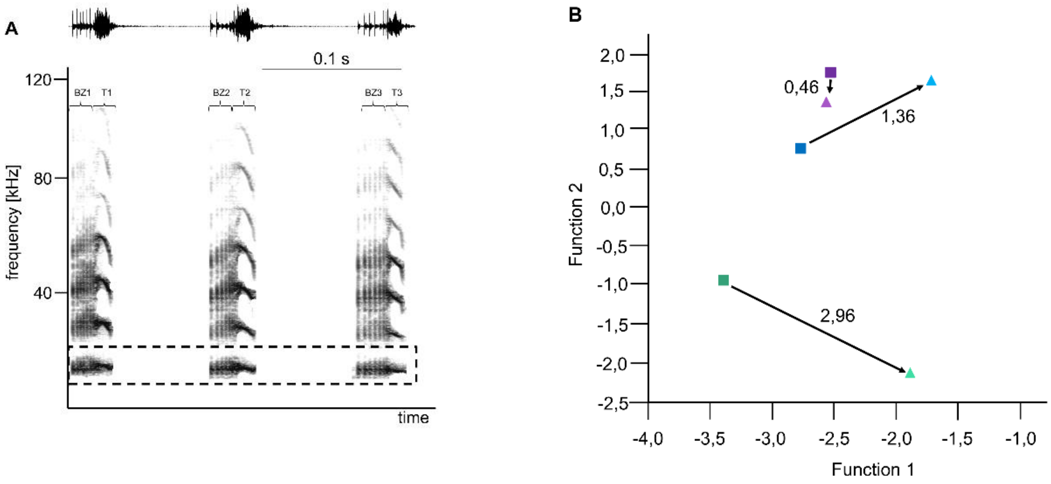


**Fig. S4. B2 syllable measurements for acoustic change tracking during ontogeny**

Fig. S4 A depicts three B2 syllables which were measured to assess acoustic change of pups during ontogeny. If possible, three syllables from three different syllable trains (i.e. sequence of at least five syllables) were selected and acoustic parameters were extracted with the software Avisoft SASLab Pro. Acoustic parameters were extracted for the pulsed part (BZ1, BZ2, BZ3 in the figure) and for the tonal part (T1, T2, T3 in the figure). Measurements were subsequently averaged (separately for pulsed and tonal part) for each train to minimize temporal dependence among syllables produced within close succession. The box illustrates that for each syllable, acoustic parameters were taken from only one harmonic containing most energy (to render comparison across syllables and syllable parts possible). As described in the method section, we used derived parameters (principal components), original parameters (e.g. duration) and LFCCs (see methods). The spectrogram was created in Avisoft SASLab Pro: Hamming window with 1024-point Fourier transformation and 87.5% overlap (frequency resolution: 244 Hz, time resolution: 0.512 ms).
Fig. S4 B shows the DFA space for one of the colonies with three exemplary pups, showing the acoustic change between the early (square) and the late (triangle) babbling phase. For each pup, we calculated individual centroids for each phase based on the DFAs. The numbers next to the arrows depict the calculated Euclidean distances. The figure includes a pup showing almost no change (violet colors), some change (blue colors) and large change (green colors).
